# Joint epitope selection and spacer design for string-of-beads vaccines

**DOI:** 10.1101/2020.04.25.060988

**Authors:** Emilio Dorigatti, Benjamin Schubert

**Affiliations:** Working group Computational Statistics, Faculty of Mathematics, Informatics and Statistics, Ludwig Maximilian Universität, München, Germany; Institute of Computational Biology, Helmholtz Zentrum München – German Research Center for Environmental Health, Neuherberg, Germany; Department of Mathematics, Technical University of Munich, Garching bei München, Germany

## Abstract

**Motivation:** Conceptually, epitope-based vaccine design poses two distinct problems: (1) selecting the best epitopes eliciting the strongest possible immune response, and (2) arranging and linking the selected epitopes through short spacer sequences to string-of-beads vaccines so as to increase the recovery likelihood of each epitope during antigen processing. Current state-of-the-art approaches solve this design problem sequentially. Consequently, such approaches are unable to capture the inter-dependencies between the two design steps, usually emphasizing theoretical immunogenicity over correct vaccine processing and resulting in vaccines with less effective immunogencity.

**Results:** In this work, we present a computational approach based on linear programming that solves both design steps simultaneously, allowing to weigh the selection of a set of epitopes that have great immunogenic potential against their assembly into a string-of-beads construct that provides a high chance of recovery. We conducted Monte-Carlo cleavage simulations to show that, indeed, a fixed set of epitopes often cannot be assembled adequately, whereas selecting epitopes to accommodate proper cleavage requirements substantially improves their recovery probability and thus the effective immunogenicity, pathogen, and population coverage of the resulting vaccines by at least two fold.

**Availability:** The software and the data analyzed are available at https://github.com/SchubertLab/JessEV

## Introduction

One of the most prominent approaches to rational vaccine design against cancer (Sahin and Türeci, 2018; Ott *et al*., 2017; Hu *et al*., 2017) and infectious diseases (Barouch *et al*., 2018; Audran *et al*., 2005) are so-called epitope-based vaccines (EVs). EVs consist of short immunogenic peptides, called epitopes, that are presented on human leukocyte antigen (HLA) molecules and elicit a T-cell response. Such vaccines can be produced quickly and cheaply with proven technologies and easily preserved. They also eliminate the risk of reversion to virulence present in regular attenuated vaccines, and can be engineered to reduce potential toxicity and inflammatory reactions (Liu, 2019).

The design process of EVs is composed of three stages: discovery of potential epitopes, selection of a subset to be included in the vaccine, and the actual design of the vaccine. As it has become clear that delivery of mixtures of separate epitopes is not effective in inducing a strong immune response, delivery strategies have been developed that assemble the selected epitopes into concatenated polypeptide vaccines, so-called string-of-beads vaccine, thereby increasing their resulting immunogenicity considerably (Yang *et al*., 1996). In a string-of-beads construct, the epitopes are linked by short sequences of few amino acids, called spacers, designed to elicit correct proteasomal cleavage at the N- and C-termini of the epitopes thereby increasing their recovery likelihood and the effective immunogencity of the vaccine.

Current state-of-the-art methods for string-of-beads design approach epitope selection and vaccine assembly independently. First, epitopes are selected to maximize the the theoretical immunogenicity of the vaccine subject to additional design constraints (Lundegaard *et al*., 2010; Toussaint *et al*., 2008) completely disregarding the assembly and processing of the vaccine. Only in a second step, the selected epitopes are assembled into a string-of-beads vaccine optimizing their recovery likelihood either using pre-determined, hand-design spacer sequences (Velders *et al*., 2001) or spacers specifically designed for each epitope pair (Schubert and Kohlbacher, 2016).

Although the immunogenicity of the selected epitopes and the cleavage likelihood of the assembled vaccine strongly influence each other (Sébastien Corneta *et al*., 2006), existing approaches cannot adequately captures and exploit this trade-off due to their sequential nature. This often leads to string-of-beads vaccines with theoretically high immunogenicity but undesirable cleavage patterns where many epitopes cannot be recovered during vaccine processing, fundamentally reducing the vaccine’s effective immunogenicity.

Our main contribution is therefore an approach that considers these two steps together using mixed integer linear programming (MILP). Our mathematical framework is able to select and assemble a subset of maximally immunogenic epitopes conforming to pre-specified design constraints regarding their conservation, coverage of pathogens and HLA alleles, as well as cleavage probabilities of their N- and C-termini and in their interior. Through extensive Monte Carlo cleavage simulations we show that the resulting vaccines provide a much greater epitope recovery rates compared to vaccines designed with a sequential approach. As a consequence, the effective immunogenicity, as well as the effective pathogen and population coverage are significantly increased, demonstrating the necessity of modeling both design steps simultaneously.

## Materials and Methods

### A unifying framework for epitope selection and assembly of string-of-bead vaccines

An epitope is effective only if it is recovered from the vaccine polypeptide (i.e., cleavage occurs at its terminals and not in its inside). Hence, epitopes should not only be selected based on their theoretical immunogencity but also based on their proteasomal cleavage likelihood and therefore their recovery probability. By controlling the cleavage likelihood through optimized arrangement of epitopes and specifically designing spacer sequences, we can increase the probability that the epitopes of a vaccine are recovered correctly. Other quantities, besides cleavage and immunogenicity, such as coverage of HLA and pathogenic variability, as well as epitope conservation might be of interest as well to make the vaccine robust and broadly applicable.

Therefore, the design problem can be described as finding the optimal set of epitopes 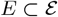 of given size *k* that maximizes the immunogencity *I*(*E*) while conforming with other pre-specified design criteria, and simultaneously assembling the epitopes into a string-of-beads vaccine with optimal spacer sequences *S_ij_* of bounded (above and below) length for each pair of connected epitopes (*e_i_, e_j_*) ∈ *E* × *E* maximizing their cleavage likelihood.

We formulate this optimization problem as a mixed integer linear program (MILP), which guarantees a global optimal string-of-beads vaccine. The MILP is conceptually divided into two blocks. The base linear program (Table 1) contains the basic constraints needed to encode the vaccine design problem, ensuring consistency of the resulting solution, reconstructing the amino acid sequence of the selected epitopes and spacers, and computing the cleavage scores for each position. The cleavage score is proportional to the cleavage probability in a specific position, and is computed as a sum of offset-dependent scores of the surrounding amino acids. As we allow spacers of variable length, it is not possible to directly calculate the offsets used to query the cleavage contributions of surrounding amino acids for a specific position in a MILP. Instead we reformulate the cleavage calculation by linearizing a bivariate function ℤ × ℤ → ℝ mapping offset and amino acid to their individual score contribution.

**Table 1:**
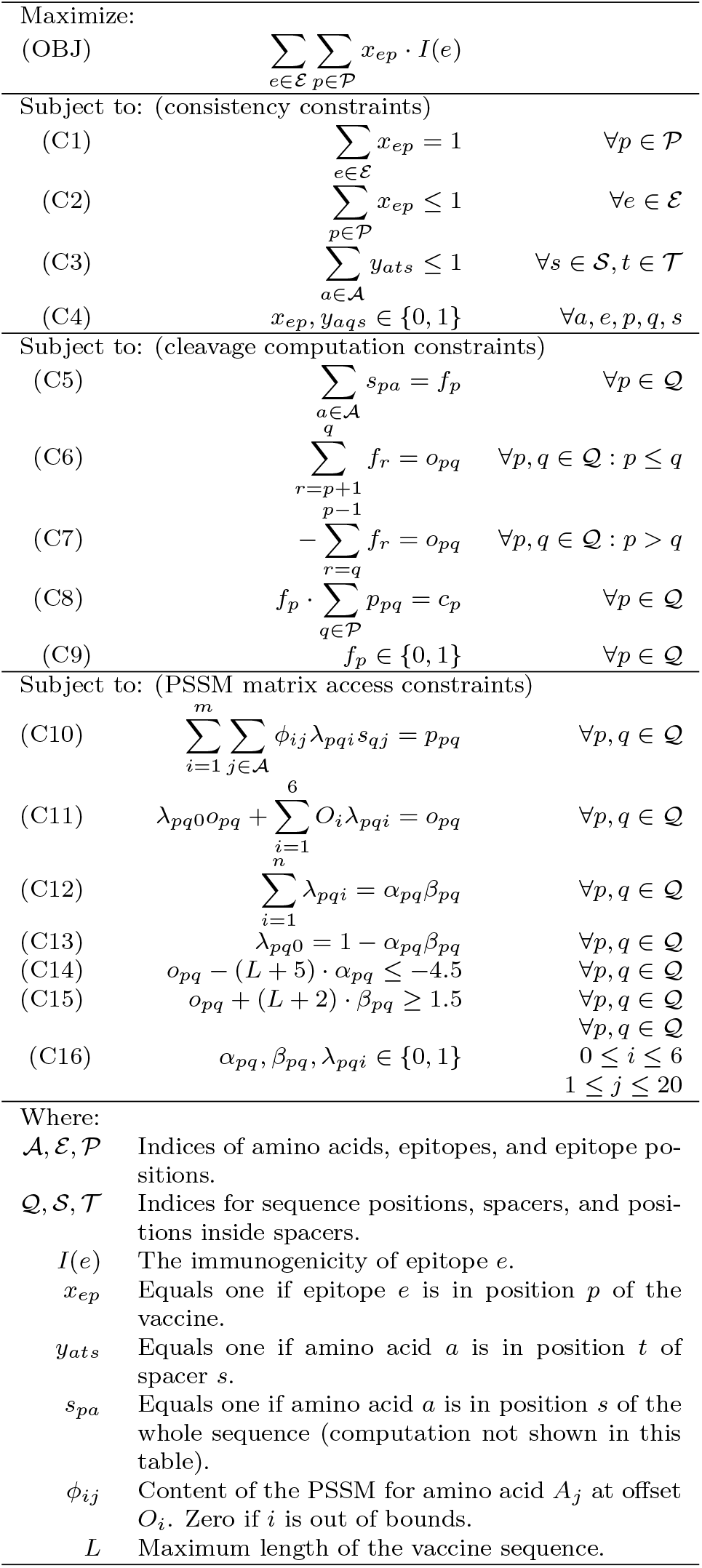
The base linear program that selects epitopes and spacers (consistency constraints), reconstructs the amino acid sequence (not shown), and computes the cleavage score for each position of the sequence (cleavage computation constraints and PSSM access constraints).

The second building block contains optional constraints related to the selection of epitopes in the vaccine and to bound cleavage scores in certain locations of the string-of-beads construct. The epitope selection constraints force the vaccine to cover a given minimum amount of pathogens and/or HLA alleles. Furthermore, they can restrict the epitopes selected to have a certain minimum average conservation. As most epitopes have extremely low conservation, we found it preferable to focus on the average, rather than the minimum. The cleavage constraints are applied to certain critical locations: the N- and C-terminals of the epitopes, their interior, and the interior of the spacers. We will later give suggestions of broadly applicable values for the cleavage site thresholds. A full description of the MILP can be found in Supplementary Table S1.

#### Immunogenicity model

As in Toussaint *et al*. (2008), we define the overall contribution of an epitope *e* to the vaccine immunogenicity as the weighted average of the log-transformed HLA binding strengths *ι_ea_* over a specified set of HLA alleles 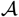:

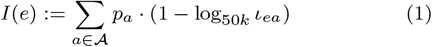

where *p_a_* is the probability of the allele *a* occurring in an individual of the target population. The set 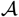 of alleles has to be chosen carefully beforehand to target a specific population. We chose HLA binding affinity as proxy of immunogenicity as it has been show to be strongly correlated with epitope immunogenicity (Sette *et al*., 1994; Paul *et al*., 2013). A variety of models have been developed predicting HLA binding affinity with high accuracy (Peters *et al*., 2020). We chose NetMHC-pan (Jurtz et al., 2017) as it is considered state-of-art and most widely used, however the framework and results presented here hold for any HLA binding affinity model.

#### Cleavage site model

Through data-driven models, we can assign a proteasomal cleavage probability to each position within a peptide sequence. State-of-art cleavage site prediction models take the amino acids of neighboring positions into account and assume their influence to be independent (Dönnes and Kohlbacher, 2005; Tenzer *et al*., 2005; Kuttler et al., 2000). Using observational data, these models estimate *p*(*A_o_* = *a*_*k*+*o*_|*C_k_* = 1), the probability that an amino acid *o* positions away from the cleavage site *k* is *a*_*k*+*o*_, and *p*(*A_o_* = *a*_*k*+*o*_), the probability of amino acid *a*_*k*+*o*_ occurring in a protein. Assuming independence, the cleavage probability is then expressed as:

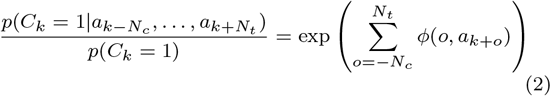

where *ϕ*(*o, a*_*k*+*o*_) is the content of a position-specific scoring matrix (PSSM) for amino acid *a*_*k*+*o*_ at offset o from the cleavage point *k*, and represents the log-ratio of the probabilities which are multiplied in Eq. 2. These model implicitly assume that *C*_*k*+*N_t_*_ = … = *C*_*k*+*N_t_*_ = 0, therefore we complement Eq. 2 with the following additional condition:

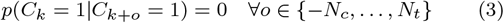

Note that the resulting score is still relative to the prior probability *p*(*C_k_* = 1) = *p_c_* of cleavage, which may vary according to the host organism. Given that the average length of the peptides cleaved by the proteasome is between seven and nine amino acids (Nussbaum *et al*., 1998), a reasonable value for this prior probability could be between 0.15 and 0.20, but as it is not clear how to set this parameter, we will investigate the influence of the prior probability ranging from zero to one. Here, we applied PCM, a PSSM proposed by Dönes and Kohlbacher that uses four C-terminal amino acids (*N_c_* = 4) and two N-terminal amino acids (*N_t_* = 1) to predict a cleavage site. It has been shown to give robust and generalizable predictions (Dönnes and Kohlbacher, 2005).

Given this model, the recovery event of an epitope *e* can then be simply computed as:

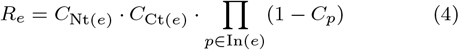

where Nt(*e*) and Ct(*e*) are the positions of *e*’s terminals, and In(*e*) are the residues inside *e*.

#### Linearizing PSSM Indexing

The difficulty in querying a PSSM within the specified MILP arises from the necessity of dynamically calculating the indexing position due to the variable length of each spacer sequence. We solve this issue by bounding the spacer length from above and below, which gives us a fixed reference frame in which we can specify the amino acid sequence of the spacer while allowing some position to be empty.

Formally, the cleavage score *Cp* at position p can be computed as follows:

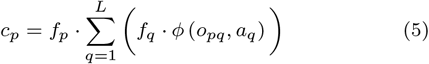

where *f_k_* = 1 if there is an amino acid in position *k, L* is the maximum length of the vaccine sequence, *ϕ*(*o, a*) is the entry of the PSSM, *a_q_* is the amino acid in position *q*, and *O_pq_* is the number of amino acids between positions *p* and *q* including the one at *q*. Eq. 5 corresponds to constraint C8 in Table 1.

The position-specific indicators *f* can be computed easily from the position-amino acid indicators (Table 1 C5) and the offset *op_q_* can be computed as the sum of the indicators for the positions between *p* (not included) and *q* (included). As *op_q_* is used to index the PSSM, we define *op_q_* to be negative when q < p and use two constraints for the positive and negative cases (Table 1 C6 and C7 respectively).

Indexing into the PSSM can then be expressed as:

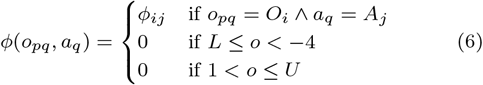

with the pivots *O_i_* ∈ {−4,…, 1} and *A_j_* ∈ {1,…, 20} indicating the offset and the amino acid, and *ϕ_j_* the entry of the PSSM. We also require to specify a lower *L* and upper bound *U* for *o*; the maximum length of the sequence suffices. All the quantities involved in Eq. 6 are integers, except for *ϕ_ij_* which is a real number.

Eq. 6 can be linearized similarly to how piece-wise linear functions are (Vielma *et al*., 2010), with a few adaptations for our specific case. In particular, we associate an indicator variable to each pivot, *s_qj_* = **1**[*a_q_* = *A_j_*] and λ_*pqi*_ = **1**[*o_pq_* = *O_i_*], and retrieve the cleavage score from the PSSM as a linear combination of these indicators with the respective pivot in constraint C10 (Table 1).

The indicators *s_qj_* can be computed easily from *x* and *y* (Supplementary Table S1). The appropriate λ can be computed by comparing every pivot *O_i_* to the actual offset *op_q_* (Table 1 C11), but require a default value of zero if *op_q_* is out of the bounds of the PSSM (i.e., −4 and 1). To this end, we introduce a new indicator λ_*pq*0_ that is not used to compute cleavage. Constraint C11 (Table 1) can always be satisfied by choosing λ_*pq*0_ = 1, therefore further constraints were added to force this to happen only if the offset is actually out of the PSSM bounds. Consequently, we introduce two additional indicator variables *α_pq_* = **1**[*o_pq_* > 1] (Table 1 C14) and *β_pq_* = **1**[*op_q_* < −4] (Table 1 C15), and set λ_*pq*0_ = 1 − *α_pq_β_pq_* (Table 1 C13).

#### Monte Carlo simulations

Given the cleavage scores, we are interested in estimating the probability that an epitope *e* of the string-of-beads vaccine is recovered, which happens when *C*_Nt_(*e*) = *C*_Ct_(*e*) = 1 and *C_p_* = 0 for *p* ∈ In(*e*) (i.e., cleavage happens at *e*’s terminals and not inside it). Eq. 2 defines the probability of cleavage at a certain position conditioned on the surrounding amino acids. To calculate *C_k_* for each position within the string-of-beads vaccine, we assume that the proteasome cleaves the vaccine from N- to C-terminus and define *p*(*C_k_* = 1|*a*_*k*−4_,…, *a*_*k*+1_) = 0 if any of *C*_*k*−4_,…, *C*_*k*−1_ have been cleaved before (i.e., ∃*C*_*k*−*i*_ = 1, *i* ∈ [1,4]). In this case we assume that *C_k_* = 0 instead. Based on this, we can define the cleavage event as a stochastic process indexed by the position *k* in the sequence:

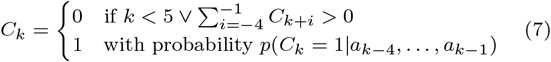

We can now estimate the recovery probability of each epitope *p*(*R_e_* = 1) by sampling from this stochastic process through Monte Carlo simulations and computing the ratio of successful recoveries, as defined in Eq. 4, over the number of simulations performed.

Using the epitope recovery probability *p*(*R_e_* = 1), we can then estimate quantities of interest such as the effective immunogenicity and effective coverage of the vaccine as the expectation of the respective metric under the recovery probability of each epitope in the vaccine. For example, the effective immunogenicity is computed as:

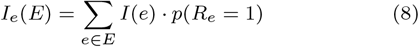

We use these simulations to evaluate the vaccines after they have been designed by solving the linear program.

### Dataset

Ninemer epitopes were extracted from 275 randomly selected sequences of the Nef gene of HIV-1 subtypes B and C, downloaded from the HIV Sequence Database (Foley *et al*., 2018; Los Alamos National Laboratory, 2019), for a total of 13,668 epitopes. We considered the same 27 HLA alleles and their frequencies as Toussaint *et al*. (2011), that together provide a theoretical coverage of 91.3% of the world population.

### Implementation

The software was implemented in Python (van Rossum, 2001), using Pyomo (Hart *et al*., 2011, 2017) to formulate the linear program and Gurobi (Gurobi Optimization, 2020) to solve it. The implementation of OptiTope (Toussaint *et al*., 2008) for epitope selection and OptiVac (Schubert and Kohlbacher, 2016) for spacer design of the sequential approach was provided by FRED2 (Schubert et al., 2016). NumPy (van der Walt et al., 2011), Scipy (Virtanen *et al*., 2020), Pandas (McKinney, 2010), statsmodels (Seabold and Perktold, 2010), Matplotlib (Hunter, 2007) and Seaborn (Waskom *et al*., 2017) were used to analyze and visualize the results in the IPython environment (Pérez and Granger, 2007)

## Results

Our vaccines are compared against a vaccine designed by first selecting the optimal set of epitopes using OptiTope (Toussaint *et al*., 2008) and then finding the optimal ordering and spacer sequences using OptiVac (Schubert and Kohlbacher, 2016), while our approach considers all 13,668 epitopes at once when designing the string-of-beads vaccine. We refer to this procedure as the *sequential* approach/design. In contrast, the method we proposed is referred to as *simultaneous* approach/design. Due to the large number of experiments required, we limited the length of all vaccines to five epitopes and at most four-amino acids spacers.

### Smaller cleavage likelihood inside epitopes and larger cleavage likelihood at their terminals is possible

We created 30 sets of 5,000 epitopes extracted without replacement from the complete set of 13,668 epitopes and designed a vaccine for each set with both sequential and simultaneous approaches.

Both approaches can reach similar cleavage likelihoods at the epitope junction sites (Fig. 2), yet the sequentially designed vaccines often exhibit less favorable cleavage patterns within epitopes (Fig. 2a). As the sequential epitope selection method cannot consider vaccine processing during epitope selection, the subsequent epitope assembly model has limited opportunity to generate a favorable cleavage pattern. Even though only 4% of epitope residues have larger cleavage than a terminal residue, 44% of them have a score larger than zero, (i.e., cleavage more likely than the prior). This leads to frequent cleavage within epitopes, which nullfies their therapeutic effected *in vivo*.

**Figure 1:**
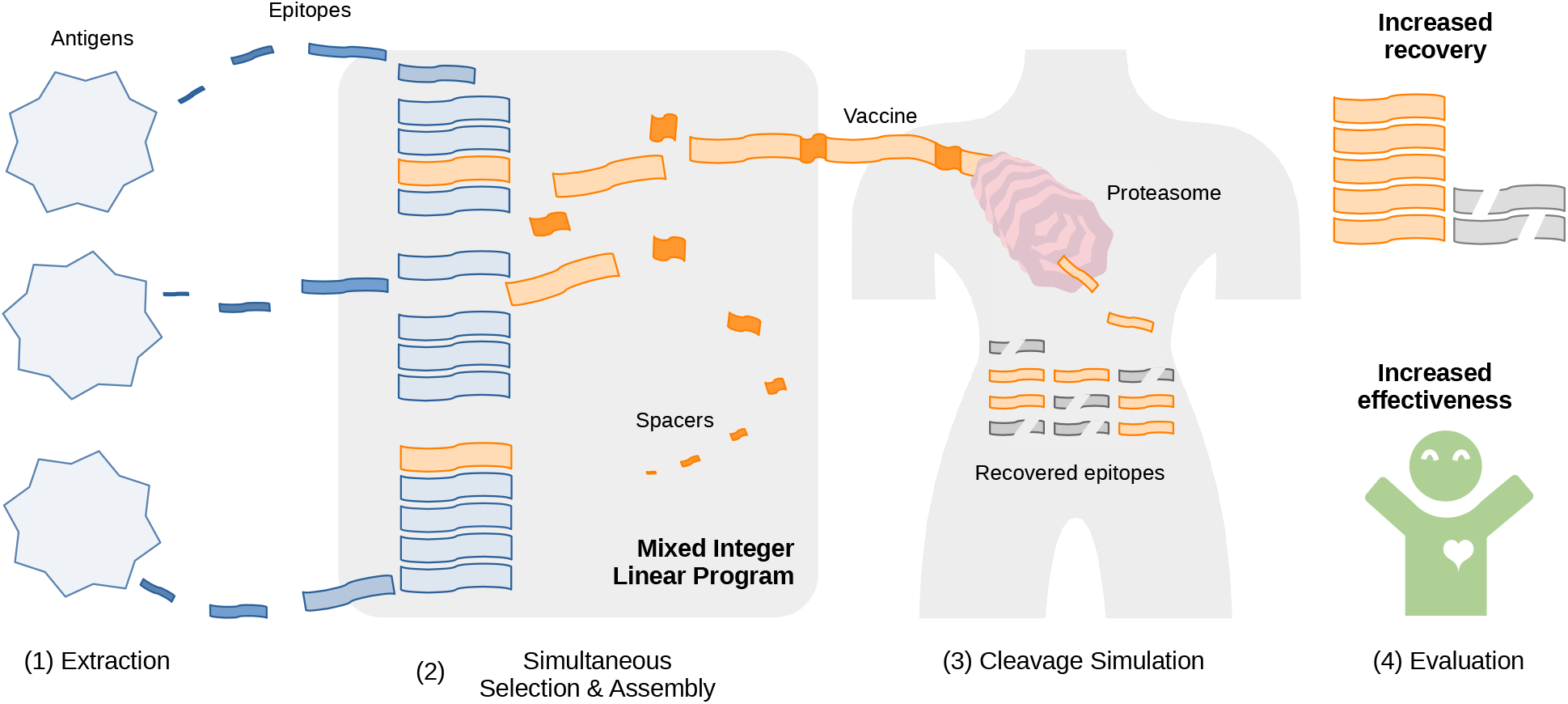
Conceptual steps in epitope-based vaccine design with the proposed framework. (1) Epitopes are extracted from a given set of antigens, and properties such as immunogenicity, coverage and conservation are computed. (2) We formulate a mixed integer linear program that creates a string-of-beads vaccine by simultaneously selecting which epitopes to include and assembling them into the final construct. This formulation maximizes the immunogenicity of the selected epitopes subject to constraints related to patient and pathogen coverage of the resulting vaccine and cleavage probability of specific residues. To connect the selected epitopes, spacers are designed to provide a high change of cleavage at the terminals of the epitopes. The vaccine will be subject to proteolytic digestion, which has strong effects on its efficacy. To quantify these effects, (3) we perform repeated stochastic simulations of proteasomal cleavage and estimate the probability that each epitope is correctly recovered from the string-of-beads construct. (4) Based on the recovered epitopes, the vaccine is evaluated in terms of the average immunogenicity of the recovered epitopes, as well as coverage and conservation with respect to the original antigens and/or the target population. We show that approaching the selection and assembly together increases the number of epitopes correctly recovered from the vaccine, making the vaccine itself more effective.

**Figure 2:**
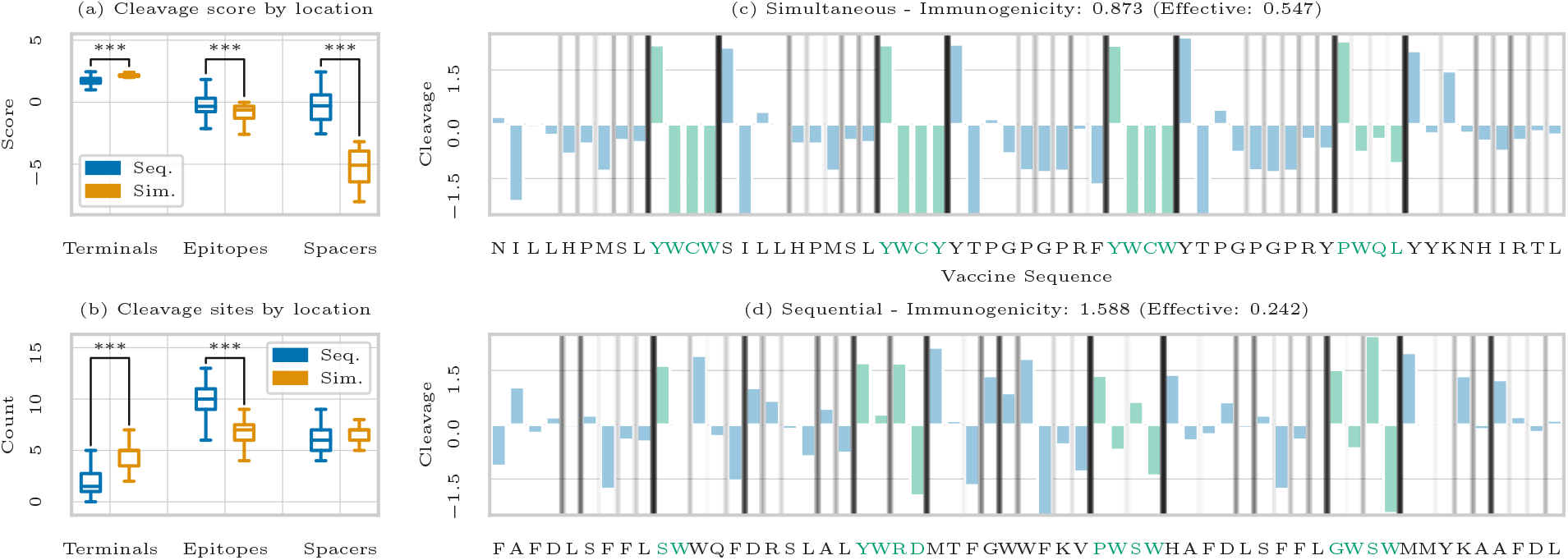
Cleavage score comparison of sequential and simultaneous approaches. (a) Shows the cleavage scores of residues at the terminals, inside the epitopes, and inside the spacers for thirty vaccines designed on random subsets of 5000 epitopes. We are able to enforce a strict separation between the scores of residues inside the epitopes and at the terminals, with a clear gap between the cleavage scores at the terminals and inside the epitopes. (b) Shows how many cleavage events, as predicted by NetChop, happened at the terminals, inside the epitopes, and inside the spacers in the same bootstraps used in (a). The marked differences are statistically significant (∗ ∗ ∗ < 0.001). (c) Shows the cleavage scores for each residue of a string-of-beads vaccine designed on the complete set of epitopes with a sequential approach and (d) with our simultaneous approach. The spacers are highlighted in green, and the gray vertical lines represent cleavage frequencies as computed by Monte Carlo simulations with a prior of 0.1, with darker shades being more likely. The title reports both theoretical and effective immunogenicity. Thanks to higher minimum cleavage at the terminals and lower maximum cleavage inside the epitopes and spacers, the effective immunogenicity of our vaccine is about twice that of the sequential approach, even though the individual epitopes are less immunogenic.

With our framework, we can take this into account and enforce negative cleavage score inside the epitopes, scores larger than two at the terminals, and smaller than negative two inside the spacers. This caused the average score of residues inside the epitopes to decrease significantly *(t* = −9.40, p-value = 9 × 10^−21^) from −0.40 for the sequential design to −1.01 for the simultaneous designs, with an effect size of −0.50. Similarly, the average scores at the terminals significantly increased (t = 16.36, p-value =1 × 10^−47^) from 1.70 to 2.16, for an effect size of 1.17. The largest difference was inside the spacers, where the average score decreased from −0.25 to −5.22 (*t* = −47.74, p-value =9 × 10^−220^, effect size of-3.62).

We then used NetChop Cterm (Nielsen *et al*., 2005) to obtain an independent prediction of the cleavage sites for every bootstrap, counting how many cleavage events happened in the spacers, terminals, and epitopes for each bootstrap (Fig. 2b). On average, there were 6.07, 1.73, and 9.40 cleavage events for the sequential design, and 6.44, 4.37, and 6.81 for the simultaneous design. The effect sizes were 0.37, 1.90 and −1.99 respectively. We then fitted a Poisson regression model considering the choice of algorithmic design as the independent variable, finding that the difference between cleavage events inside the spacers was not significant (0.06 ± 1.06, *z* = 0.57 and *p*-value = 0.57), but the difference in terminals and epitopes were (0.92±0.17, *z* = 5.56, *p*-value = 3×10^−8^ and −0.32±0.09, *z* = −3.39, *p*-value = 7 × 10^−4^).

The bounds we imposed on the cleavage scores forced the solver to pick epitopes with lower immunogenicity, but thanks to the improved recovery rates the effective immunogenicity was 0.10 ± 0.15 larger than for the sequential approach (*t* = 6.69, *p*-value = 1 × 10^−8^ for an effect size of 1.86). However, due to this restriction on the available epitopes, the problem proved infeasible in three cases out of 30. Relaxing the bounds on the cleavage scores could prevent infeasibility.

Comparing vaccines designed with the same procedure on the complete set of epitopes reveals that the epitopes selected in the simultaneous solution have a combined immunogenicity of 0.87 (Fig. 2c), only 55% of the immunogenicity of the sequential approach (Fig. 2d), but the resulting effective immunogenicity was 0.54, 240% larger than that of the sequential approach (using a prior cleavage probability of 0.1). For this and all subsequent experiments, we relaxed the negative cleavage score constraint on the first three residues after the N-terminal of the epitopes. Under our cleavage model, these three residues cannot be cleaved when the N-terminal is, therefore their score is uninfluential as long as the N-terminal is cleaved frequently. This resulted in a 79% increase in immunogenicity and 68% increase in effective immunogencity.

### Increased epitope recovery rates improve effective immunogenicity and coverage

We designed string-of-bead vaccines using several thresholds for minimum termini cleavage *ν* and *γ* (ranging from 1.5 to 2.5) and maximum epitopes’ interior cleavage score thresholds *η* (from −1 to 1). To optimize for effective coverage, we additionally performed a grid search on pathogen conservation (from 5% to 20%) while keeping pathogen coverage at 99% and allowing larger values of *η* (between 1 and 2). We then performed 1,000 Monte Carlo cleavage simulations using different prior cleavage probabilities p*c*, and selected the values for *ν*, *γ*, and *η* that resulted in the largest average effective immunogenicity or coverage for each *p_c_*. Finally, we compared the best simultaneous solution with the fixed vaccine produced by the sequential approach (Fig. 3).

**Figure 3:**
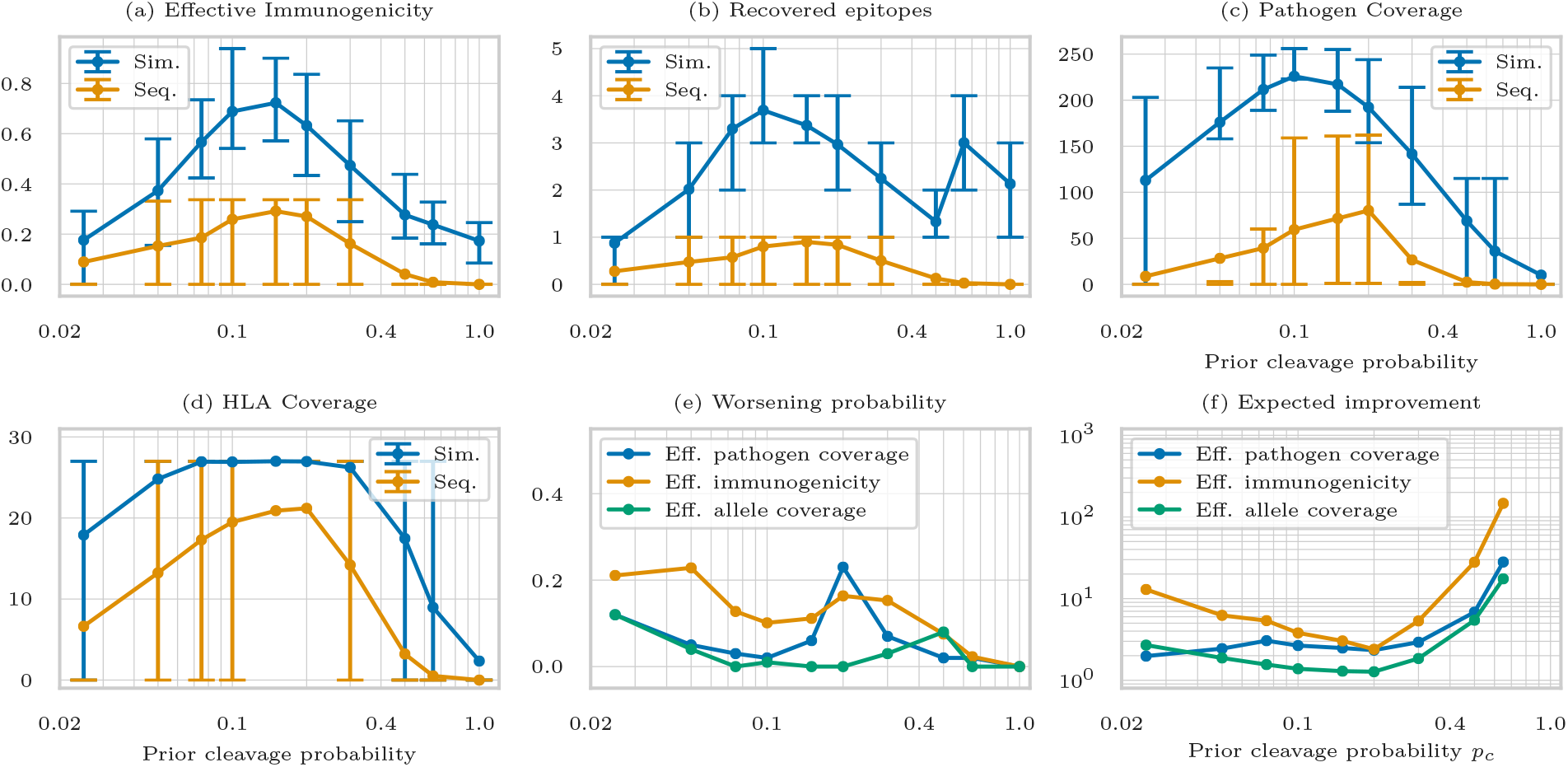
Evaluation of string-of-beads designed with our simultaneous approach and a sequential approach across different prior cleavage probabilities. Figures **(a), (b), (c)**, and **(d)** show mean, 25-th and 75-th percentile of the Monte Carlo simulations for effective immunogenicity, recovered epitopes, pathogen coverage, and HLA coverage respectively. Our solution is better under all metrics across all choices of prior cleavage probabilities. Note that both vaccines optimize for immunogenicity in figures (a) and (b), and for coverage in figure (c) and (d), which means that different constraints are used to produce them. Figures **(e)** and **(f)** show the probability of worsening and expected improvement of effective immunogenicity, effective pathogen coverage, and effective HLA coverage. Both were estimated through five thousand bootstrap of the outcomes of the thousand Monte Carlo simulations. String-of-bead vaccines produced by our simultaneous approach are very frequently not worse than the sequential approach, and on average between three to five times better across a realistic range of prior probabilities. At cleavage probabilities larger than 0.7, no epitopes are ever recovered for the sequential approach, hence the expected improvement approaches infinity.

The effective immunogenicity of vaccines designed with the simultaneous method was consistently larger by at least 97% than that resulting from a sequential designs across all settings of *p_c_*; often, the 25-th percentile of was larger than the 75-th percentile of the sequential design (Fig. 3a). The number of recovered epitopes was also at least 315% larger (Fig. 3b). However, the higher epitope recovery rates were partly offset by the lower immunogenicity of the selected epitopes, as forcing low cleavage likelihoods inside of the selected epitopes restricted the set of candidates. A qualitatively similar result could be observed for effective pathogen (Fig. 3c) and HLA coverage (Fig. 3d). The simultaneous string-of-beads designs outperformed the sequentially designs by a margin of at least 140% and 27% for effective pathogen and allele coverage respectively. We were able to design vaccines such that 2.13 epitopes were recovered on average even with *p_c_* = 1 by enforcing the interior epitope cleavage score to be smaller than −1, corresponding to a cleavage probability below *e*^−1^ ≈ 0.37. In practice, however, half of the residues inside the epitopes of such a vaccine had a cleavage score smaller than −3.91 (*e*^−3,91^ ≈ 0.02), showing that favorable cleavage can be achieved even in the most adverse conditions.

We then bootstrapped the Monte Carlo trials to quantify the probability that simultaneous designs had worse immunogenicity and HLA/pathogen coverage across different prior recovery probabilities. On average, they were worse 12%, 7%, and 1% of the times for effective immunogenicity, pathogen coverage and HLA coverage respectively (Fig. 3e). Given the low number of HLA alleles, there was a significant probability that both methods could cover the same number of alleles (62% on average, and 42% that both effectively covered zero alleles). For pathogens this probability was lower but still considerable (average 24%), while the effective immunogenicity was equal only 5% of the times.

We also quantified the expected improvement for each *p_c_* (i.e., the ratio between the average effective immunogenicity of the two solutions) and found that simultaneous designs consistently outperformed the state of the art by two- to three-fold, with greater improvement as *p_c_* approached one (Fig. 3f).

### The same constraints are effective across a realistic range of prior cleavage probabilities

Different settings of *ν, γ*, and *η* were needed to obtain the largest possible effective immunogenicity depending on the prior cleavage probability *p_c_*. As *p_c_* increased, *ν, γ*, and *η* decreased, as the smaller cleavage score was offset by the larger prior probability. Fig. 4c traces the evolution of these parameters and shows that for 0.15 ≤ *p_c_* < 0.5 the optimal settings were *ν* = *γ* = 1.95 and *η* = −0.1. A prior between 0.15 and 0.20 resulted in fragments of length between 7 and 10 on our dataset, consistent with what was observed *in vitro* (Nussbaum *et al*., 1998), suggesting that these are promising values for experimental testing. Additionally, Fig. 4a shows that for these prior probabilities the effective immunogenicity of the second, third, and fourth best settings was within 5% of the best design for that prior, meaning that the effective immunogenicity for these priors was not overly sensitive to the settings of *ν*, 7 and *η*.

**Figure 4:**
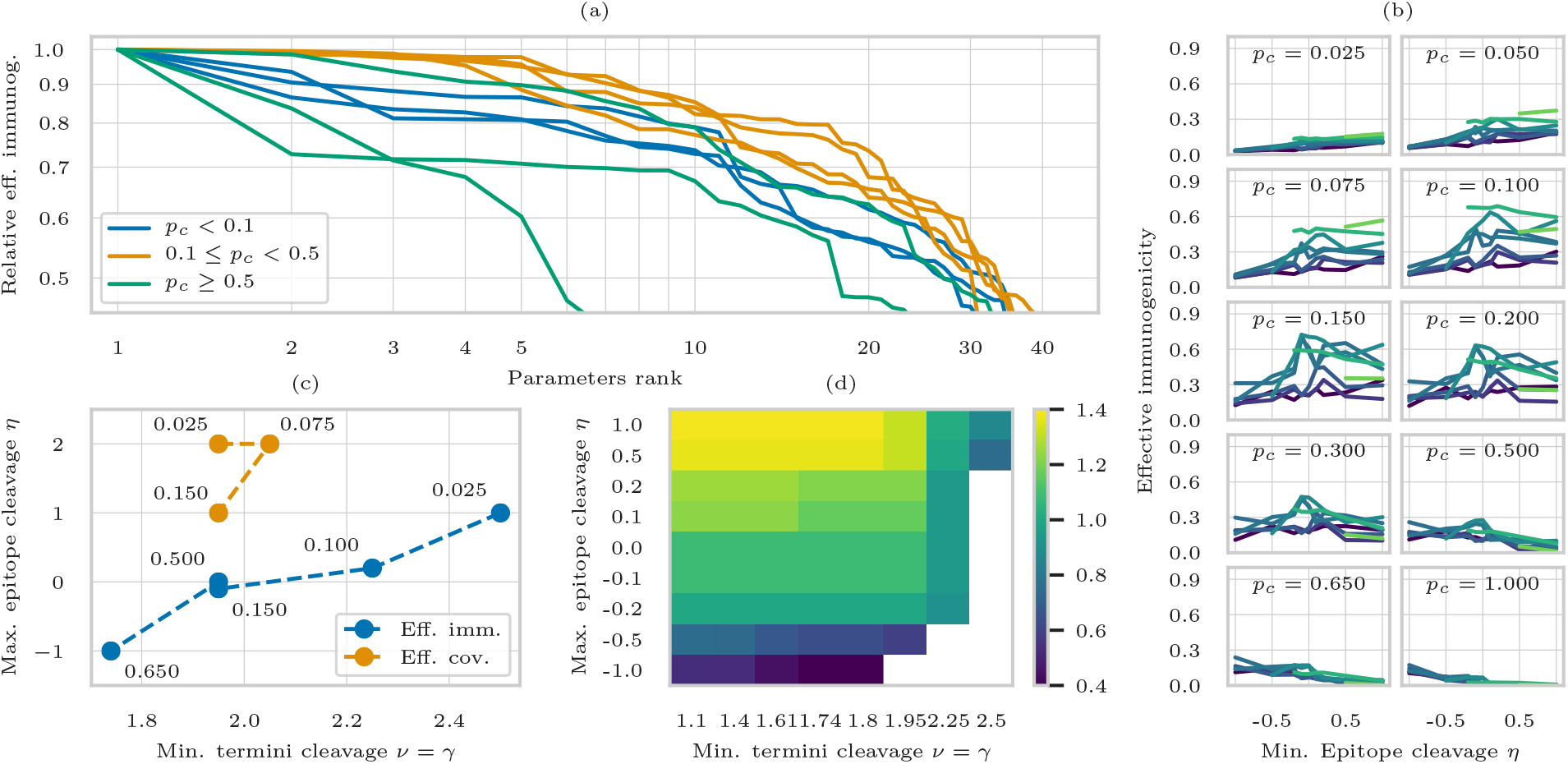
The effects of cleavage constraints on the immunogenicity objective and effective immunogenicity. (a) for each prior probability, we plot the effective immunogenicity relative to the best obtained for that prior (*y* axis) for different parameter settings ranked by effective immunogenicity (*x* axis). There is a range of prior probabilities, from 0.1 to 0.3, where four or five different parameter settings are within 5% of the largest effective immunogenicity. (b) effective immunogenicity (*y*-axis) as a function of the inner epitope cleavage (*x*-axis) for different cleavage at the termini (lighter lines for larger constraints) and nine different prior cleavage probabilities (in each sub-figure). For prior cleavage probabilities in a reasonable range, the best effective immunogenicity is obtained with an inner epitope cleavage around zero, while lower settings work best for high priors and larger ones for low priors. (c) the effect of prior cleavage probability (annotated close to each data point) on parameters (*x* and *y* axes) that result in the largest effective immunogenicity (blue) or effective pathogen coverage (orange). Only transitions are displayed, meaning that several prior probabilities between, for example, 0.15 and 0.5 (not included) have the same optimal settings for the effective immunogenicity. As the prior cleavage probability increases, constraints on cleavage at the termini can be relaxed, while the score inside the epitopes must be kept lower. Optimizing for effective coverage requires larger possible cleavage likelihoods inside the epitopes, but similar cleavage likelihoods at the termini. (d) immunogenicity objective for different cleavage constraints, with light background for infeasible settings. Enforcing low cleavage likelihoods inside the epitopes greatly reduces the immunogenicity objective, as many epitopes are not eligible due to higher cleavage likelihoods in the residues of their second half, which cannot be reduced through the preceding spacer.

In general, *η* was more critical than *ν* and 7, since it had a considerable effect on the set of epitopes that could be selected (Fig. 4d). In fact, according to our cleavage model, a spacer affects cleavage only in the first four residues of the following epitope, while the score in the following five residues cannot be altered. For prior cleavage probabilities in a reasonable range, the largest effective immunogenicity was obtained with *η* close to zero, whereas larger absolute values caused a reduction in the effective immunogencity. For very high (low) prior probabilities, the best results were obtained with low (high) values for *η* (Fig. 4b).

This also explained why optimizing for effective coverage required larger values for *η* as very few epitopes were conserved across a sufficient number of pathogens and were excluded when *η* was too small. In fact, the epitopes with the highest pathogenic coverage raking in the top 1%, 2% and 5% percentile covered 21%, 13% and 5% of the pathogenic antigens respectively. This illustrates that including conserved epitopes is fundamental even if they are recovered less frequently.

### Optimized spacers are necessary but few variants are used

Inspecting the vaccines with largest effective immunogenicity for each of the 10 prior cleavage probabilities revealed that only nine different spacer sequences were used. Two sequences, MWQW and MWRW, were used in 19 out of 40 spacers. These spacer sequences increased the C-terminal cleavage score by 2.37 and 2.32 respectively, corresponding to a 10-fold likelihood increase. Only five of the possible 20^4^ sequences induce a larger increase. However, they all end in K, which reduces the N-terminal score by 1.4 and greatly limits the number of viable epitopes after the spacer.

Designing vaccines with fixed MWQW spacer results in a reduction in immunogenicity, both theoretical and effective, of 35-45% compared to the optimized spacers. Similarly, using the popular spacer AAY causes a decrease of 85-95%, and required to relax the cleavage constraints on *ν* and *γ* from 1.95 to 1.6, and *η* from 0 to 1.0. This highlights the need for spacers designed *ad hoc*.

## Discussion and Conclusions

No current state-of-art design approach is able to simultaneously select epitopes and assemble them into a string-of-beads vaccine construct. Our work fills this gap through a linear programming formulation that guarantees optimality of the design. This linear program finds a set of epitopes of maximal immunogenicity, as well as their arrangement and spacers linking them, ensuring that the vaccine simultaneously satisfies constraints related to pathogen, HLA coverage, conservation, and cleavage likelihood in critical positions of the construct.

Being based on mixed-integer linear programming renders the simultaneous epitope selection and assembly problem of string-of-beads vaccines NP-hard. In most cases, this does not prevent the solver to find a solution in a reasonable time, in part because of the many heuristics (Fischetti and Lodi, 2011; Bixby *et al*., 2000; Berthold, 2006) that can be employed. Though, certain constraint configurations can make the solving process slower. In these rare cases most time is usually spent on improving a solution whose objective value is already within at most a few percent of the optimum. As this gap is known during the process, the solver can be interrupted early, obtaining an almost-optimal solution with formal guarantees on its quality. Indexing the position-specific scoring matrix to compute cleavage scores contributes a great deal to the overall complexity of the linear program, as its size in terms of variables and constraints scales quadratically with the maximum number of residues in the vaccine. Using spacers of fixed length can therefore significantly reduce the computational resources needed to find a vaccine, at the price of slightly longer polypeptides.

We assumed a simple stochastic model of proteasomal cleavage and used Monte Carlo simulations to estimate the recovery probability of each epitope in a vaccine to show that approaching the epitope selection and epitope assembly problems together results in increased recovery probability of the vaccine’s epitopes. We also verified our results with NetChop Cterm (Nielsen *et al*., 2005), an independent proteasomal cleavage prediction tool, to confirm that, in spite of the simplistic nature of the cleavage predictor we used in the linear program, our approach significantly reduces cleavage sites inside the epitopes and increases cleavage frequency at the terminals. This, in turn, translated to improved effective immunogenicity, coverage, and conservation. The main reason for this improvement can be traced to the ability of our framework to select epitopes that have a small cleavage probability in their interior, thus preventing unwanted cleavage in these locations. We also argued that this constraint should be relaxed in order to include highly conserved epitopes, as the gains in coverage offset the reduction in recovery frequency.

Our framework depends on the ability to express the computation of the cleavage score in a linear form. Non-linear functions can be approximated by piece-wise linear approximations, and lookup tables can be used in the worst case, but this could make solving even small instances of the linear program impractical due to extremely long runtimes. As a precaution against the possibility that our cleavage model is too simplistic, stricter cleavage constraints than what our simultations suggest could be enforced. Moreover, finding the right bounds for the cleavage scores requires an outer optimization loop where the vaccines are evaluated using the Monte Carlo simulations. In this work we performed a grid search to study the influence of the parameters on the effective immunogenicity, but more complicated optimization strategies such as Bayesian optimization (Brochu *et al*., 2010; Shahriari *et al*., 2016) can be used to reduce the computational requirements needed to find solutions with good effective immunogenicity or coverage.

In conclusion, our approach allows precise control of the cleavage probability of every residue in a string-of-beads construct through simultaneously approaching epitope selection and vaccine assembly. This allows us to greatly improve the recovery probability of the epitopes in the construct, which translates to increased effectiveness of the vaccine as a whole.

