## Supplementary Table S1 for "Joint epitope selection and spacer design for string-of-beads vaccines"

### Supplement One - Complete MILP Formulation

| | | |
|---|---|---|
| **Maximize:** | | |
| (OBJ) | $\sum_{e\in\mathcal{E}}\sum_{p\in\mathcal{Q}} x_{ep}\cdot I(e)$ | |
| **Subject to: (consistency constraints)** | | |
| (C1) | $\sum_{e\in\mathcal{E}} x_{ep} = 1$ | $\forall p \in \mathcal{P}$ |
| (C2) | $\sum_{p\in\mathcal{Q}} x_{ep} \le 1$ | $\forall e \in \mathcal{E}$ |
| (C3) | $\sum_{a\in\mathcal{A}} y_{ats} \le 1$ | $\forall s \in \mathcal{S}, t \in \mathcal{T}$ |
| (C4) | $x_{ep}, y_{aqs} \in \{0,1\}$ | $\forall a, e, p, q, s$ |
| **Subject to: (cleavage computation constraints)** | | |
| (C5) | $\sum_{a\in\mathcal{A}} s_{pa} = f_p$ | $\forall p \in \mathcal{Q}$ |
| (C6) | $\sum_{r=p+1}^{q} f_r = o_{pq}$ | $\forall p,q \in \mathcal{Q} : p \le q$ |
| (C7) | $-\sum_{r=q}^{p-1} f_r = o_{pq}$ | $\forall p,q \in \mathcal{Q} : p > q$ |
| (C8) | $f_p \cdot \sum_{q\in\mathcal{P}} p_{pq} = c_p$ | $\forall p \in \mathcal{Q}$ |
| (C9) | $f_p \in \{0,1\}$ | $\forall p \in \mathcal{Q}$ |
| **Subject to: (PSSM matrix access constraints)** | | |
| (C10) | $\sum_{i=1}^{m}\sum_{j\in\mathcal{A}} \phi_{ij}\lambda_{pqi}s_{qj} = p_{pq}$ | $\forall p,q \in \mathcal{Q}$ |
| (C11) | $\lambda_{pq0}o_{pq} + \sum_{i=1}^{6} O_i\lambda_{pqi} = o_{pq}$ | $\forall p,q \in \mathcal{Q}$ |
| (C12) | $\sum_{i=1}^{n} \lambda_{pqi} = \alpha_{pq}\beta_{pq}$ | $\forall p,q \in \mathcal{Q}$ |
| (C13) | $\lambda_{pq0} = 1 - \alpha_{pq}\beta_{pq}$ | $\forall p,q \in \mathcal{Q}$ |
| (C14) | $o_{pq} - (L+5)\cdot\alpha_{pq} \le -4.5$ | $\forall p,q \in \mathcal{Q}$ |
| (C15) | $o_{pq} + (L+2)\cdot\beta_{pq} \ge 1.5$ | $\forall p,q \in \mathcal{Q}$ |
| (C16) | $\alpha_{pq}, \beta_{pq}, \lambda_{pqi} \in \{0,1\}$ | $\forall p,q \in \mathcal{Q}$ <br> $0 \le i \le 6$ <br> $1 \le j \le 20$ |
| **Subject to: (Epitope selection constraints)** | | |
| (C17) | $\sum_{e\in\mathcal{E}}\sum_{p\in\mathcal{Q}} x_{ep}\tau_{eo} \ge \theta_o$ | $\forall o \in \mathcal{O}$ |
| (C18) | $\sum_{o\in\mathcal{O}} \theta_o \ge \Theta$ | |
| (C19) | $\sum_{e\in\mathcal{E}}\sum_{p\in\mathcal{P}}\sum_{o\in\mathcal{O}}\left[x_{ep}\left(\sum_{o\in\mathcal{O}}\tau_{eo} - \Gamma\right)\right] \ge 0$ | |
| (C20) | $\theta_o \in \{0,1\}$ | $\forall o \in \mathcal{O}$ |
| **Subject to: (Cleavage constraints)** | | |
| (C21) | $\sigma \le i_{s(\bar{e}+\bar{s})+\bar{e}+t}$ | $\forall s \in \mathcal{S}, t \in \mathcal{T} : t > 1$ |
| (C22) | $c_{s(\bar{e}+\bar{s})+\bar{e}+t} \le \Sigma$ | $\forall s \in \mathcal{S}, t \in \mathcal{T}$ |
| (C23) | $c_{e(\bar{e}+\bar{s})+o} \ge \eta$ | $\forall e \in \mathcal{E}$ <br> $1+\epsilon < o < \bar{e}$ |
| (C24) | $c_{p(\bar{e}+\bar{s})} \ge \nu$ | $\forall p \in \mathcal{Q} : p > 0$ |
| (C25) | $c_{s(\bar{e}+\bar{s})+\bar{e}} \ge \gamma$ | $\forall s \in \mathcal{S}$ |
| **Subject to: (sequence reconstruction constraints)** | | |
| (C26) | $\sum_{e\in\mathcal{E}\text{ s.t. } e[o(p)]=a} x_{e,s(p)} \le s_{pa}$ | $\forall p \in \mathcal{P} : o(p) < E$ <br> $a \in \mathcal{A}$ |
| (C27) | $\sum_{a\in\mathcal{A}} s_{pa} \le 1$ | $\forall p \in \mathcal{Q} : o(p) < E$ |
| (C28) | $s_{pa} \ge y_{s(p),o(p)-\bar{e},a}$ | $\forall p \in \mathcal{Q} : o(p) \ge E$ <br> $a \in \mathcal{A}$ |
| (C29) | $\sum_{a\in\mathcal{A}} s_{pa} \le \sum_{a\in\mathcal{A}} y_{s(p),o(p)-\bar{e},a}$ | $\forall p \in \mathcal{Q} : o(p) \ge E$ |
| (C30) | $s_{pa} \in \{0,1\}$ | $\forall p \in \mathcal{Q}, a \in \mathcal{A}$ |

| **Where:** | |
|---|---|
| $\mathcal{A}, \mathcal{E}, \mathcal{P}$ | Indices of amino acids, epitopes, and epitope positions. |
| $\mathcal{Q}, \mathcal{S}, \mathcal{T}$ | Indices of sequence positions, spacers, and positions inside spacers. |
| $x_{ep}$ | One if epitope $e$ is in position $p$. |
| $y_{aqs}$ | One if position $q$ of spacer $s$ contains amino acid $a$. |
| $I(e)$ | The immunogenicity of epitope $e$. |
| $e[o]$ | The amino acid in position $o$ of epitope $e$. |
| $s_{pa}$ | One if amino acid $a$ is in position $p$ of the whole sequence. |
| $f_p$ | One if there is an amino acid in position $p$. |
| $o_{pq}$ | How many amino acids are between positions $p$ and $q$. Negative if $q < p$. |
| $p_{pq}$ | The cleavage contribution of the amino acid in position $q$ when computing cleavage for position $p$. It is zero if $p$ is empty. |
| $O_i$ | $i$-th offset used to index the PSSM matrix (between -4 and 1, inclusive). |
| $\phi_{ij}$ | Content of the PSSM matrix at offset $O_i$ and amino acid $A_j$. |
| $\lambda_{pqi}$ | One if the offset $o_{pq}$ is $O_i$. |
| $\lambda_{pq0}$ | One if $o_{pq}$ is not in the bounds of the PSSM matrix. |
| $\alpha_{pq}$ | One if $-L \le o_{pq} < -4$. |
| $\beta_{pq}$ | One if $1 < o_{pq} \le L$. |
| $L$ | Maximum length of the vaccine sequence. |
| $\mathcal{O}$ | Set of possible options with respect to which compute coverage and/or conservation. |
| $\tau_{eo}$ | One if epitope $e$ covers option $o$. |
| $\Theta$ | Minimum coverage. |
| $\Gamma$ | Minimum average conservation. |
| $c_p$ | Cleavage score at position $p$. |
| $\sigma$ | Minimum cleavage inside spacers. |
| $\Sigma$ | Maximum cleavage inside spacers. |
| $\eta$ | Maximum cleavage inside epitopes. |
| $\epsilon$ | Cleavage inside epitopes is not enforced to the first $\epsilon$ positions. |
| $\nu$ | Minimum cleavage at the N-terminus. |
| $\gamma$ | Minimum cleavage at the C-terminus. |
| $\bar{e}, \bar{s}$ | Length of epitopes and spacers. |
| $s(p)$ | The *segment* that $p$ belongs in, $s(p) = \lfloor p/(\bar{e}+\bar{s})\rfloor$, with $\bar{e}$ the length of the epitopes, and $\bar{s}$ the maximum spacer length. |
| $o(p)$ | The *offset* within the segment that $p$ belongs in,$o(p) = p \bmod (\bar{e}+\bar{s})$. |

Table S1: Complete MILP
